## Supplementary Material for "Neuronal tuning and population representations of shape and category in human visual cortex"

**Single channel analysis – LFP results:** Out of the 332 visually responsive LFP sites, 79 sites were significantly selective for the shape type dimension alone, whereas 19 were category – selective, and 100 had interactions between shape type and category (chi2 = 66.8, $p < 0.00001$). Note that a small number of channels (N = 1 and N=2 in array 2 and 3, respectively) did not show any main effect or interaction effect in the 2-way ANOVA, but were nevertheless stimulus-selective based on a 1-way ANOVA on the net responses to all 54 stimuli.

**Decoding – LFP results:** We obtained similar results when decoding shape type and category using the high-gamma responses (Fig S8A), although a few differences compared to the MUA decoding were noticeable. Array 2 and 3 exhibited reliable category decoding over a time window of several hundreds of ms (which was shorter than for the MUA decoding), while category decoding on array 1 barely reached significance (and was sustained for the MUA). Array 4 also barely reached statistical significance for both factors due to the low signal-to-noise ratio of the LFP on this array.

**Figure S1: Selectivity Index**

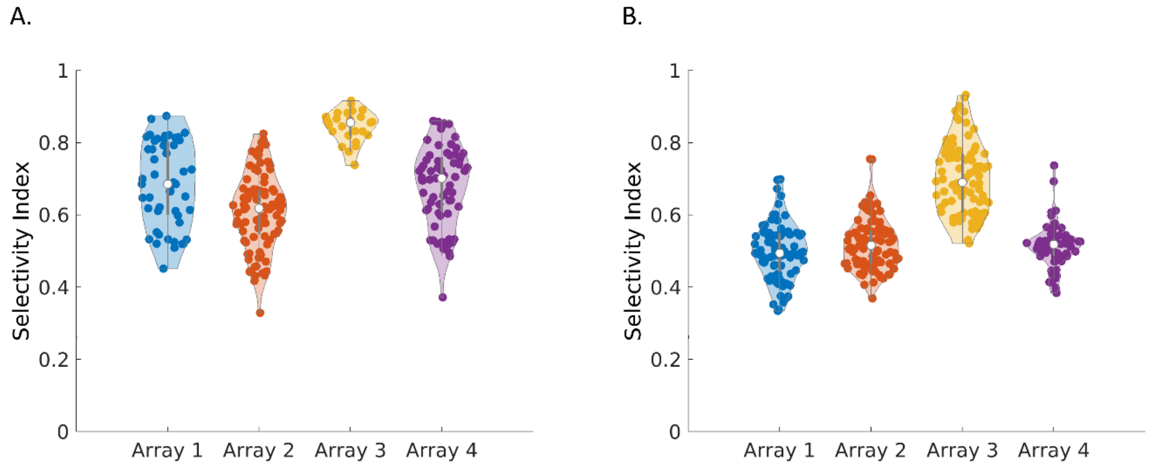

**Figure S2: Eta$^2$**

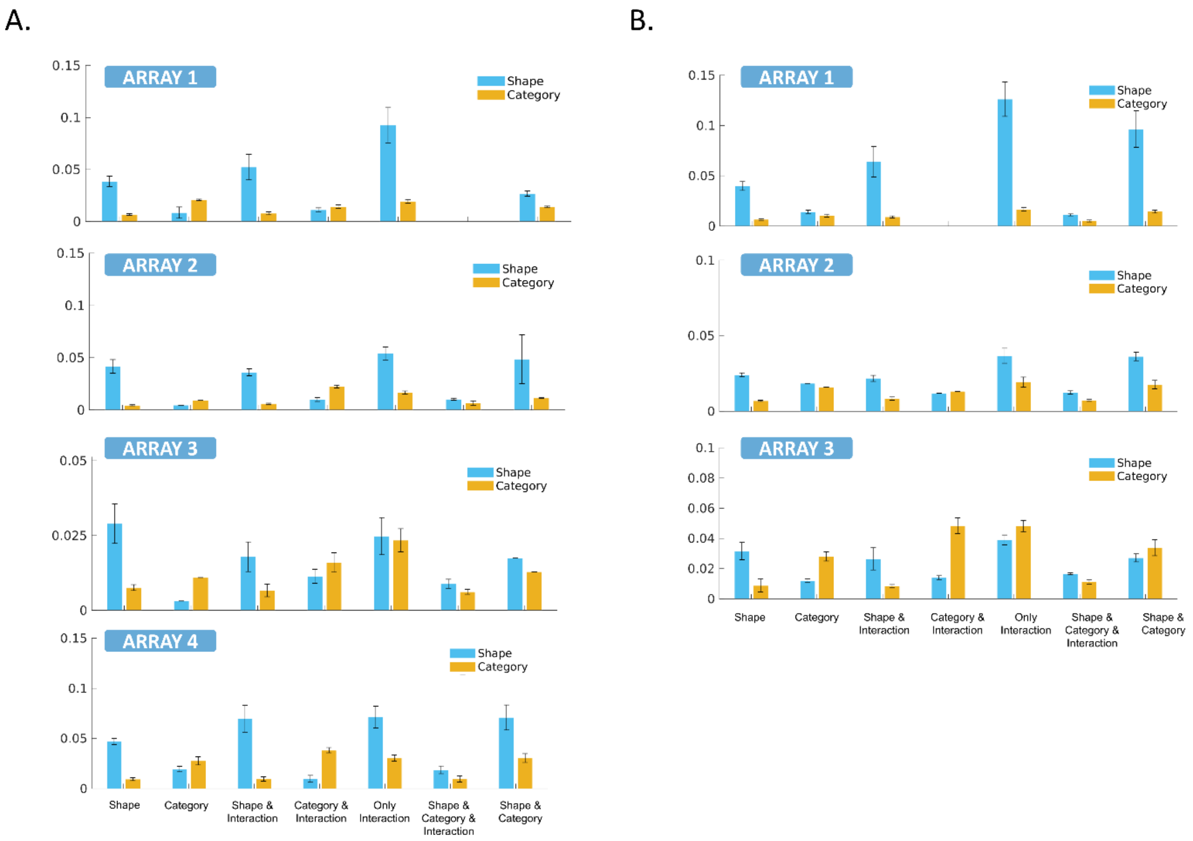


**Figure S3: Dissimilarity analysis for the high – gamma responses**

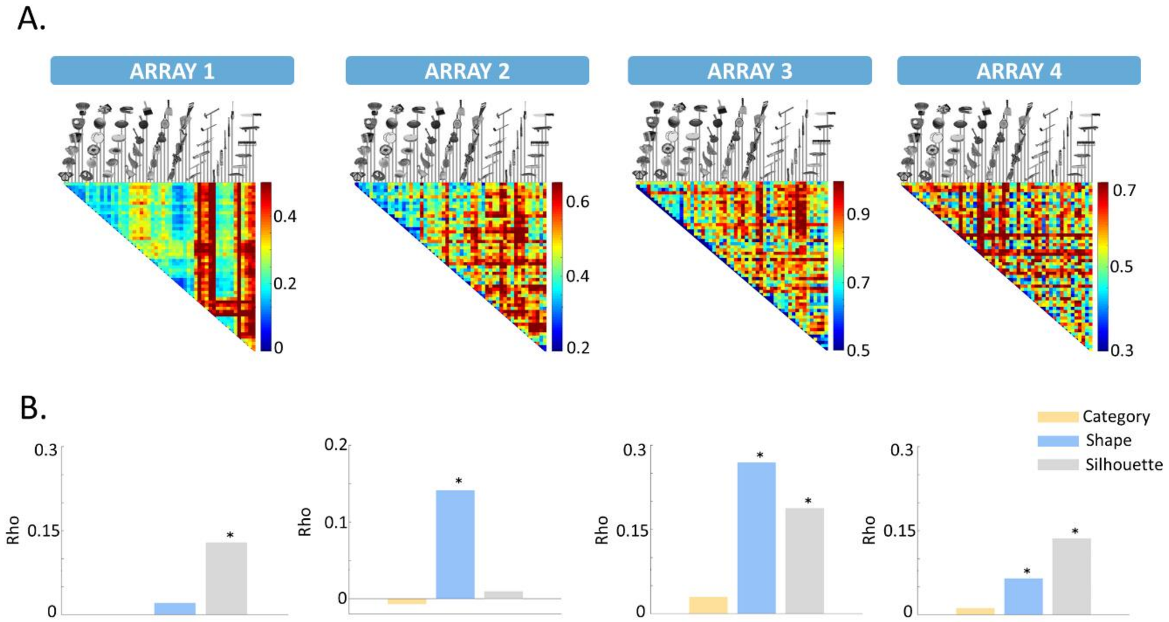

**Figure S4: Multidimensional scaling for the high – gamma neural dissimilarity matrices**

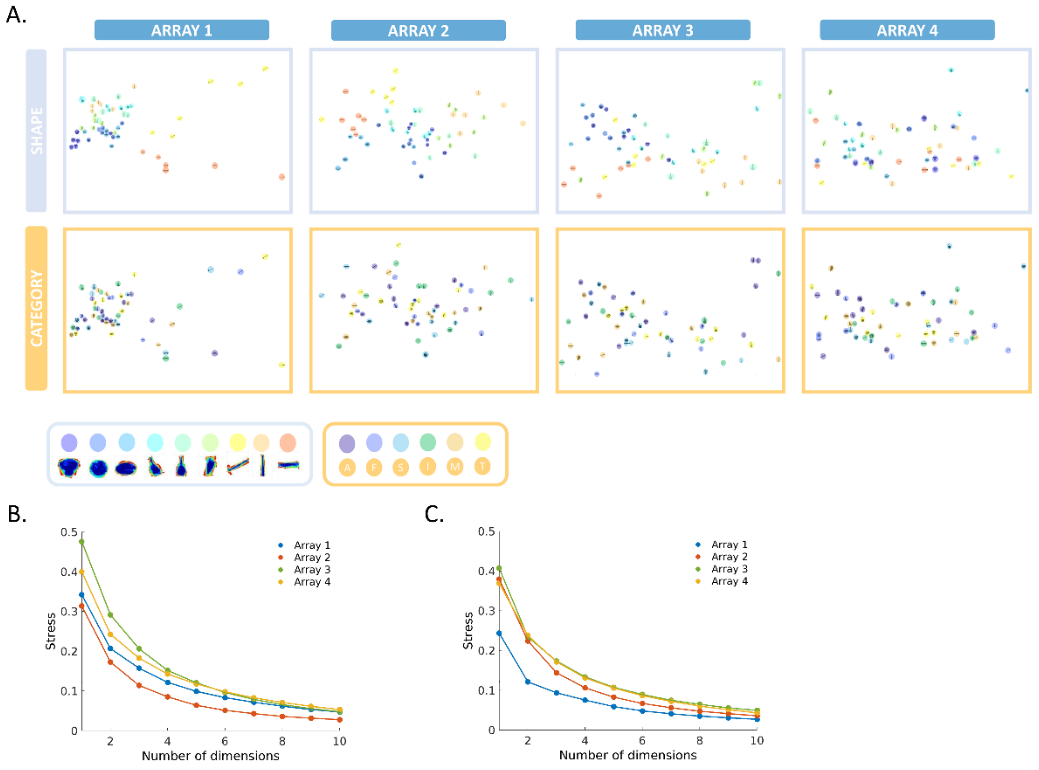


**Figure S5: Hierarchical cluster analysis**

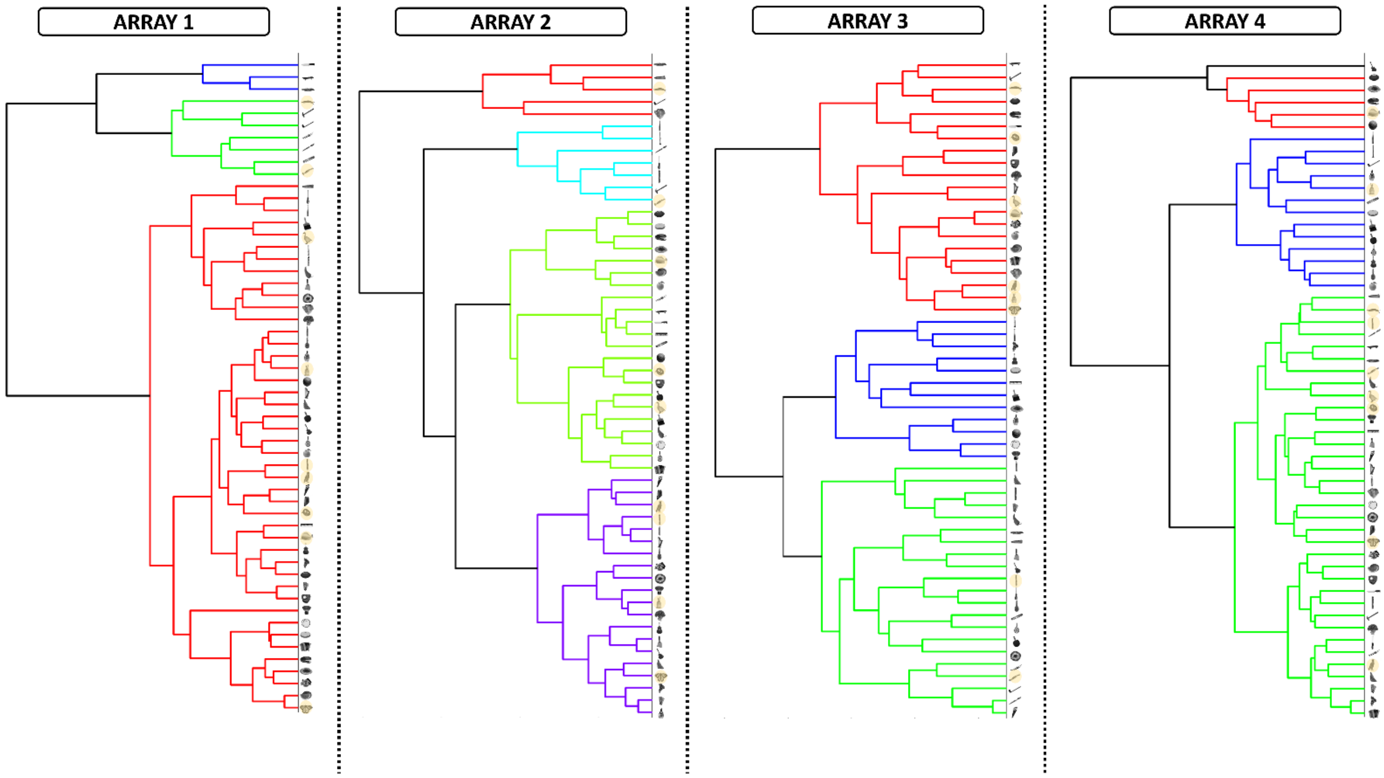

### Figure S6: Linear decoding of the high – gamma responses

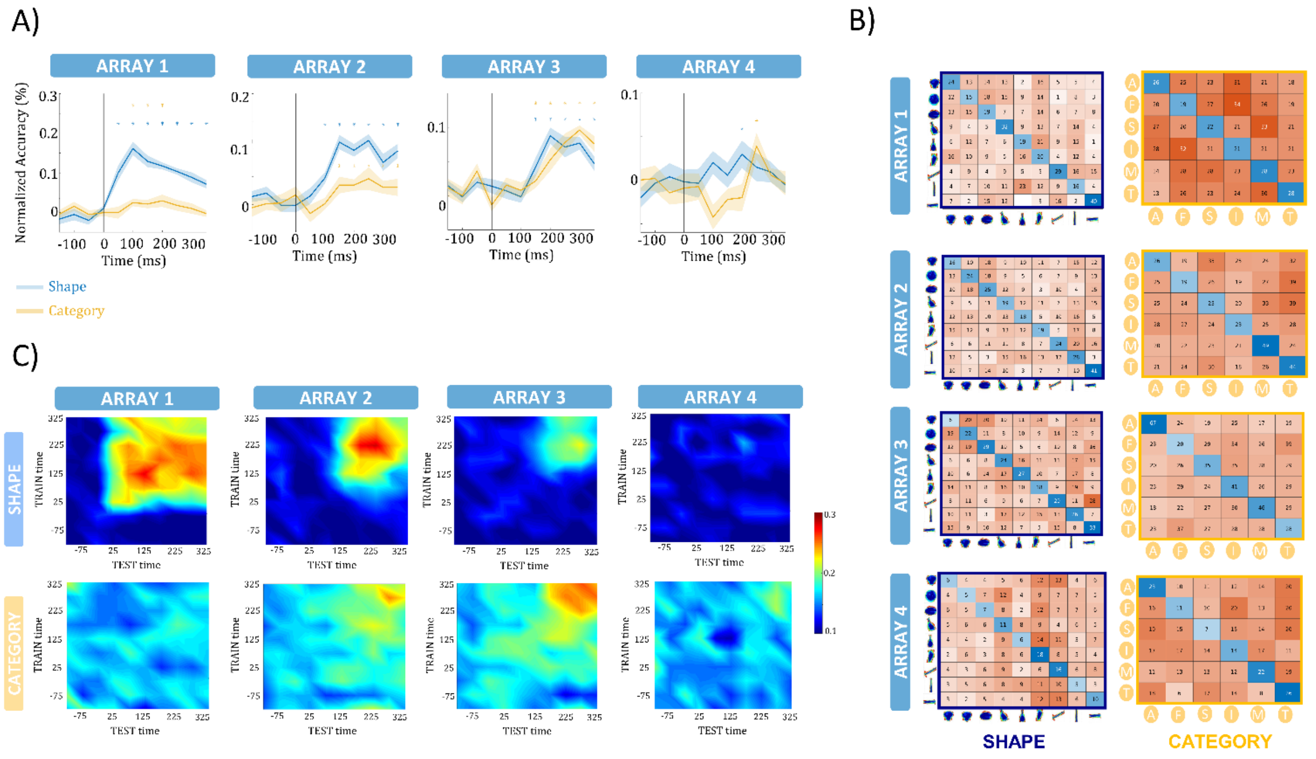


### Figure S7: Decoding accuracy after removing the class “Animals”

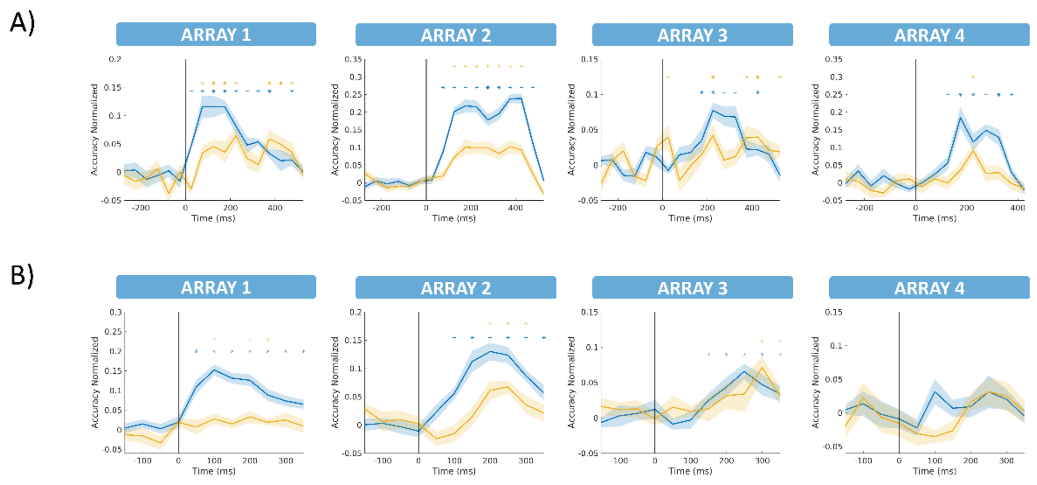

**Table S1: Summary of the 2 – way ANOVA statistics for the example sites in Figure 2.**

| Example site | SS | df | MS | F | $eta^2$ | p – value |
|---|---|---|---|---|---|---|
| $MUA_1$ - Shape | 19237 | 8 | 2404.6 | 6.34 | 0.05 | 0.000 |
| $MUA_1$ - Category | 1978 | 5 | 395.6 | 1.04 | 0.005 | 0.52 |
| $MUA_1$ - Interaction | 16827 | 40 | 420.7 | 1.11 | 0.04 | 0.65 |
| $MUA_2$ - Shape | 3421 | 8 | 427.7 | 1.86 | 0.014 | 0.06 |
| $MUA_2$ - Category | 2832 | 5 | 566.5 | 2.46 | 0.011 | 0.02 |
| $MUA_2$ - Interaction | 17036 | 40 | 425.9 | 1.85 | 0.069 | 0.0007 |
| $MUA_3$ - Shape | 84939 | 8 | 10617 | 4.43 | 0.03 | 0.00003 |
| $MUA_3$ - Category | 11171 | 5 | 2234 | 0.93 | 0.004 | 0.46 |
| $MUA_3$ - Interaction | 322515 | 40 | 8062 | 3.36 | 012 | 0.000 |
| $LFP_1$ - Shape | 1215167 | 8 | 151895.9 | 6.07 | 0.06 | 0.000 |
| $LFP_1$ - Category | 191338 | 5 | 38267.7 | 1.53 | 0.009 | 0.17 |
| $LFP_1$ - Interaction | 1280152 | 40 | 32003.8 | 1.28 | 0.06 | 0.11 |
| $LFP_2$ - Shape | 1577496 | 8 | 197187.1 | 8.8 | 0.05 | 0.16 |
| $LFP_2$ - Category | 2310706 | 5 | 462141.3 | 20.6 | 0.07 | 0.000 |
| $LFP_2$ - Interaction | 5853641 | 40 | 146341 | 6.53 | 0.18 | 0.000 |
| $LFP_3$ - Shape | 771272 | 8 | 96409 | 4.06 | 0.04 | 0.0001 |

| | | | | | | |
|---|---|---|---|---|---|---|
| $LFP_3$ - Category | 241338 | 5 | 48267 | 2.03 | 0.01 | 0.07 |
| $LFP_3$ - Interaction | 1675346 | 40 | 41883 | 1.76 | 0.08 | 0.003 |

**Table S2: Results of Representational Similarity Analysis (RSA) conducted on the high-gamma neural dissimilarity matrices**

| **ARRAYS** | **Category** | **Shape** | **Silhouette** |
|---|---|---|---|
| 1 | Rho = -0.01, p = 0.66 | Rho = 0.02, p = 0.21 | Rho = 0.13, p = 0.00 |
| 2 | Rho = -0.01, p = 0.6 | Rho = 0.14, p = 0.00 | Rho = 0.01, p = 0.36 |
| 3 | Rho = 0.03, p = 0.15 | Rho = 0.27, p = 0.00 | Rho = 0.19, p = 0.00 |
| 4 | Rho = 0.01, p = 0.34 | Rho = 0.06, p = 0.006 | Rho = 0.14, p = 0.00 |

### LEGENDS:

**Figure S1:** Violin plots depicting the distribution of Selectivity Indices for visually responsive channels in each array, at both the MUA **(A)** and LFP **(B)** levels.

**Figure S2: A)** Average eta$^2$ values of the shape (blue bars) and the category (orange bars) dimension for all MUA sites with significant effects. The height of each bar indicates the average value across sites and the error bar represents the standard error. For arrays 1, 2, and 4, the eta$^2$ values for shape are consistently higher than those for category across all sites. In array 3, however, the difference is less pronounced for sites without a significant main effect of shape. **B)** eta$^2$ values for all high - gamma sites with arrays 1, 2, and 3 showing similar results to those observed at the MUA level. Array 4 is excluded due to a lack of channels with significant effects.

**Figure S3: A)** Neural dissimilarity matrices for all arrays based on the high - gamma responses. **B)** Results of RSA for category – similarity (orange), shape – similarity (blue), and silhouette – similarity (grey). The asterisks indicate the significance of the correlation.

**Figure S4: A)** MDS performed on the high - gamma neural dissimilarity matrices shows pairwise distances in a 2D space for each array. The 2D arrangements are color – coded first according to the 9 different shape – types (upper panel), and then according to the 6 different semantic categories (lower panel). **B)** Stress level / goodness of fit of the MDS at the MUA level for 1 to 10 dimensions. **C)** Stress level / goodness of fit of the MDS at the high - gamma level for 1 to 10 dimensions

**Figure S5:** The hierarchical plot shows the structure of the neural dissimilarity matrix of each array. Results reveal clustering according to shape type for arrays 1,2, and 4 but not for array 3. Array 3 shows more clustering according to category, specifically for the “Animal” category (orange – shaded stimuli).

**Figure S6: A)** Temporal evolution of the SVM normalized decoding accuracy for the shape (blue) and the category (orange) dimension at the high – gamma level. The shaded region around the line represents the standard error across the cross validations. The asterisks indicate the significance of the accuracy. **B)** Confusion matrices are illustrating the performance of the decoding per class for the shape (upper panel) and the category (lower panel) dimension for a specific time – window (arrays 1,2: 75 - 275 ms, array 3: 175 – 275 ms, array 4: 125 – 225 ms) at the high - gamma level. The classification performance of array 3 for the category dimension is predominantly restricted to the "animals" category. **C)** Generalization of the decoders over time for the shape (upper panel) and the category (lower panel) dimension. The y – axis corresponds to the TRAIN time window, the x – axis to the TEST time – window and the colors to the accuracy level of the decoding.

**Figure S7:** Temporal evolution of the SVM normalized decoding accuracy for the shape (blue) and the category (orange) dimension at the MUA (A) and high – gamma (B) level after removing the “Animals”

class. The shaded region around the line represents the standard error across the cross validations. The asterisks indicate the significance of the accuracy. For arrays 3 and 4 the accuracy is considerably lower and less significant without the “Animals” category.

**Table S1:** Summary of the 2 – way ANOVA statistics for the example sites in Figure 2. The columns, presented from left to right, represent the following key statistical measures ; Sum of Squares (SS): The total sum of squares associated with the ANOVA model ; Degrees of Freedom (df): The degrees of freedom associated with each source of variation in the ANOVA model ; Mean Squares (MS): The average sum of squares for each source of variation ; F-Statistic (F): The calculated F-value for each source of variation, indicating the ratio of between-group variation to within-group variation ; Eta Squared (η2): The effect size measure representing the proportion of variance explained by each source of variation ; p-Value: The significance level associated with each source of variation.

**Table S2:** Results of Representational Similarity Analysis (RSA) conducted on the high-gamma neural dissimilarity matrices. The following key measures are reported: Rho (Pearson Correlation): Rho represents the Pearson correlation coefficient, quantifying the similarity between the neural dissimilarity matrices and the behavioral dissimilarity matrices ; p: The p-value associated with the correlation coefficient, indicating the level of statistical significance.
